# Filamentous Bacteriophage Delay Healing of Pseudomonas-Infected Wounds

**DOI:** 10.1101/2020.03.10.985663

**Authors:** Michelle S. Bach, Christiaan R. de Vries, Arya Khosravi, Johanna M. Sweere, Medeea Popescu, Jonas D. Van Belleghem, Gernot Kaber, Elizabeth B. Burgener, Dan Liu, Quynh-Lam Tran, Tejas Dharmaraj, Maria Birukova, Vivekananda Sunkari, Swathi Balaji, Nandini Ghosh, Shomita S. Mathew-Steiner, Sundeep G. Keswani, Niaz Banaei, Laurence Nedelec, Chandan K. Sen, Venita Chandra, Patrick R. Secor, Gina A. Suh, Paul L. Bollyky

## Abstract

We have identified a novel role for filamentous bacteriophage in the delayed healing associated with chronic *Pseudomonas aeruginosa (Pa)* wound infections. Pf phage delays wound re-epithelialization in the absence of live *Pa,* indicative that Pf effects on wound healing are independent of *Pa* pathogenesis. Pf phage directly inhibits autocrine signaling of CXCL1 (KC) to impede keratinocyte migration and wound re-epithelization. In agreement with these studies, a prospective cohort study of 36 human patients with chronic *Pa* wound infections revealed that wounds infected with Pf positive strains of *Pa* took longer to heal and were more likely to increase in size compared to wounds infected with Pf negative strains. Together, these data implicate Pf phage in the delayed wound healing associated with *Pa* infection through direct manipulation of mammalian target cells. We propose that Pf phage may have potential as a biomarker and therapeutic target in delayed wound healing.

## Introduction

Chronic wounds are associated with extensive human suffering and massive economic costs (Järbrink et al., 2016; Kirsner, 2016). Over 6.5 million Americans have chronic wounds, and their care is estimated to cost $25 billion annually (Kirsner, 2016). Bacterial infections frequently complicate chronic wounds, leading to delayed wound healing (Guo and DiPietro, 2010), increased rates of amputation (Gardner and Frantz, 2008), and extensive morbidity and mortality (Bowler, 2002; Clinton and Carter, 2015; Scali and Kunimoto, 2013).

Wounds infected with the Gram-negative pathogen *Pseudomonas aeruginosa (Pa)* are characterized by poor outcomes (Dowd et al., 2008; James et al., 2008; Kirketerp-Møller et al., 2008; Malic et al., 2009). *Pa* is prevalent in infections of burns (Tredget et al., 2004), diabetic ulcers, and post-surgical sites (Percival et al., 2015). The presence of *Pa* is associated with delayed wound closure in both humans as well as animal models (Bryers, 2008; Guo and DiPietro, 2010; Watters et al., 2013; Zhao et al., 2012), and wounds infected with *Pa* tend to be larger than those in which *Pa* is not detected (Gjødsbøl et al., 2006; Jesaitis et al., 2003). This wound chronicity is associated with ineffective bacterial clearance (Sen et al., 2009) and delayed wound healing (Bjarnsholt et al., 2008). In addition to wound infections, *Pa* is a major human pathogen in other contexts (Bryers, 2008; Høiby et al., 2011; Sen et al., 2009) due to increased antibiotic resistance incidence. In recognition of the magnitude of this problem, *Pa* is deemed a priority pathogen by the World Health Organization (WHO) and the Centers for Disease Control (CDC) (Centers for Disease Control and, 2013; Tacconelli et al., 2018). Therefore, there is great interest in identifying novel biomarkers, virulence factors, and therapeutic targets associated with *Pa* infections.

One such potential target is Pf bacteriophage (Pf phage), a filamentous virus produced by *Pa* (Dowd et al., 2008; Malic et al., 2009; Secor et al., 2020). Unlike lytic bacteriophages used in phage therapy (Gordillo Altamirano and Barr, 2019; Górski et al., 2020), Pf typically does not lyse its bacterial hosts. Instead, Pf phage integrate into the bacterial chromosome as prophage and can be produced without destroying their bacterial hosts (Rakonjac et al., 2011; Roux et al., 2019). Indeed, the production of phage virions is partially under bacterial control (Castang and Dove, 2012). We and others have reported that Pf phage contribute to *Pa* fitness by serving as structural elements in *Pa* biofilms (Secor et al., 2015; Tarafder et al., 2020) and contributing to bacterial phenotypes associated with chronic *Pa* infection, including aggregation (Secor et al., 2018) and reduced motility (Secor et al., 2017).

Pf phage also impact host immunity in ways that promote chronic infection. We recently reported that Pf phage directly alter cytokine production and decrease phagocytosis by macrophages (Secor et al., 2017; Sweere et al., 2019). These effects were associated with Pf triggering maladaptive anti-viral immune responses that antagonize anti-bacterial immunity. Consistent with these effects, *Pa* strains that produced Pf phage (Pf(+) strains) were more likely than strains that do not produce Pf (Pf(-) strains) to establish wound infections in mice (Sweere et al., 2019). These and other studies implicate Pf phage as a virulence factor in *Pa* infections (Burgener et al., 2019; Rice et al., 2009; Secor et al., 2015).

We hypothesized that Pf phage might contribute to the delayed wound healing often associated with *Pa* wound infections. To test this, we first examined the effects of Pf phage on wound healing and barrier function in an *in vivo* mouse and pig models. We then investigated the mechanism by which Pf phage inhibits wound healing in a well-established *in vitro* model (keratinocyte scratch assay). Finally, we examined the association between Pf phage and delayed wound healing in a cohort of 36 patients seen at the Stanford Advanced Wound Care Center.

## Results

### Infection with a Pf(+) strain of *Pa* causes worse morbidity in a murine chronic wound infection model

We first examined the impact of a Pf(+) strain of *Pa* (PAO1) compared to a Pf(-) isogenic strain (PAO1ΔPf4) on morbidity and mortality in a delayed inoculation murine wound infection model. We previously pioneered this model to study the initial establishment of *Pa* wound infections in otherwise healthy C57BL/6 mice in the absence of foreign material, diabetes, or other forms of immune suppression (Sweere et al., 2019; de Vries et al., 2020). Here, we have adapted this model to study wound healing in the setting of infection over a two-week time period. In brief, we generated bilateral, full-thickness, excisional wounds on the dorsum of mice. These wounds were then inoculated 24 hours later with 4 × 10^5^/mL PAO1, PAO1ΔPf4, or PBS as a control. We then assessed wound size daily through day 13 post-*Pa* inoculation (day 14 post-wounding). A schematic of this protocol is shown in **Figure 1A**.

**Figure 1.**
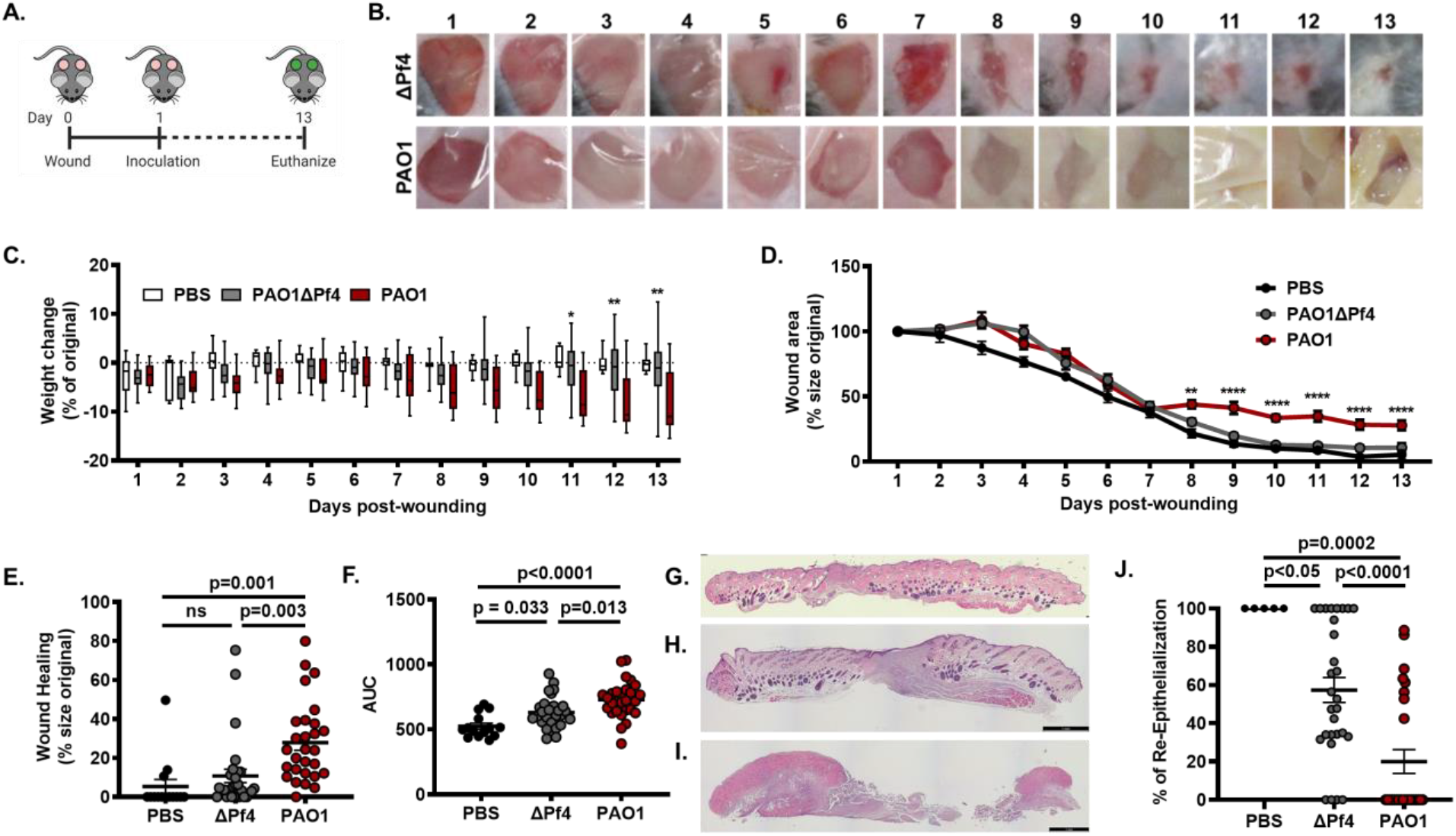
Pf phage lead to decreased wound healing in murine models. A) Schematic of the full thickness delayed-inoculation chronic *Pa* wound infection murine model. B) Images of representative mice from PAOΔPf4 and PAO1groups from Day 1-13 post-wounding. C) Weight loss of mice across all days post-wounding (For PAO1 vs PAO1ΔPf4, Day 11: p<0.05; Day 12: p<0.01; Day 13: p<0.01; two-way ANOVA). D) Wound healing rates for all wounds across all days from Day 1-13 post wounding. n=28 wounds in PAOΔPf4 and PAO1groups; n=14 wounds in PBS group. Results are mean % size of original wound ± SE (**p<0.005 ****p<0.0001 by one-way ANOVA). E) Wound healing at Day 13 post-wounding measured as percentage of original wound area. F) Area-under-curve (AUC) analysis for wound healing rates compiled for all wounds for all Days 1-13 post-wounding (n=28 wounds in PAOΔPf4 and PAO1 groups; n=14 wounds in PBS group; comparison by one-way ANOVA). G) Representative H&E stain of Day 13 uninfected PBS wound. H) Representative H&E stain of Day 13 wound infected with PAO1ΔPf4. I) Representative H&E stain of Day 13 wound infected with PAO1. J) Percent re-epithelialization calculated as ((Day 1 epithelial gap – Day 13 epithelial gap)/Day 1 epithelial gap). Statistics by one-way ANOVA).

All mice survived the 14-day experiment (**Figure S1**) and all wounds were infected on Day 3 (**Figure 1B**), regardless of whether they received PAO1 or PAO1ΔPf4. However, by day 13 wounds in the PAO1 group remained inflamed and purulent. In contrast, PAO1ΔPf4 wounds were largely healed (**Figure 1B)**. Moreover, mice inoculated with PAO1 lost significant weight compared to mice that received PAO1ΔPf4 or PBS (**Figure 1C**). These data confirm Pf phage contributes to worse morbidity in this chronic wound model, consistent with prior studies showing Pf promotes *Pa* pathogenesis (Secor et al., 2020; Sweere et al., 2019).

### Pf4 phage delay wound healing

We next examined the effect of Pf4, the phage present in PAO1, on wound healing in our *in vivo* model. Healing was defined as the remaining open wound area as a fraction of the original wound area. Inoculation with PAO1 ΔPf4 led to significantly improved wound healing over time compared to infection with PAO1. Indeed, wound healing in many of the mice inoculated with PAO1 plateaued by day 7 (**Figure 1D)**. By Day 13, most of the mice from the PAO1ΔPf4 group resolved their infections and healed their wounds completely, whereas most wounds infected with PAO1 persisted (**Figure 1E**). Similar results were observed when area under curve (AUC), a measure of cumulative change in wound area, was evaluated (**Figure 1F)**. We then examined wound histology from tissue sections collected on Day 13 post-wounding that were collected from these mice. We observed that uninfected wounds and wounds infected with PAO1ΔPf4 exhibited healthy granulation tissue **(Figure 1G-H)** whereas wounds infected with PAO1 had substantial necrotic debris, minimal granulation tissue, and obvious infection (**Figure 1I**). When the epithelial gap was measured, wounds infected with PAO1 were found to have significantly less re-epithelialization by day 13 compared to PAO1ΔPf4 **(Figure 1J)**.

Taken together, these data indicate that the production of Pf4 phage by PAO1 is associated with increased morbidity, impaired wound healing, and delayed re-epithelialization in this model.

### Pf4 phage delay re-epithelialization in the absence of bacterial infection

Because the aforementioned data were collected in the setting of infection, it is difficult to distinguish wound persistence is due to phage-enhanced *Pa* pathogenesis (Sweere et al., 2019) or from a direct inhibition on wound healing by Pf. To elaborate if Pf directly inhibits wound healing, we treated mice with purified Pf4 phage in the absence of bacterial infection. In particular, we administered heat killed PAO1 (HK-*Pa*) or PBS to C57BL6 mice one day after wounding in the absence or presence of supplemental Pf4 phage. Day 7 rather than Day 13 was used in the studies as wounds healed more rapidly in the absence of live bacterial infections. In addition, wounds were splinted with a silicone ring to prevent healing via contraction which otherwise happens in mice in the absence of infection (Wang et al., 2013). A schematic detailing this protocol is shown in **Figure 2A**. Both the wound area on Day 1 (**Figure 2B**) and the epithelial gap on Day 7 post-wounding (**Figure 2C**) were assessed morphometrically. Representative area demarcations are shown inset into these images.

**Figure 2.**
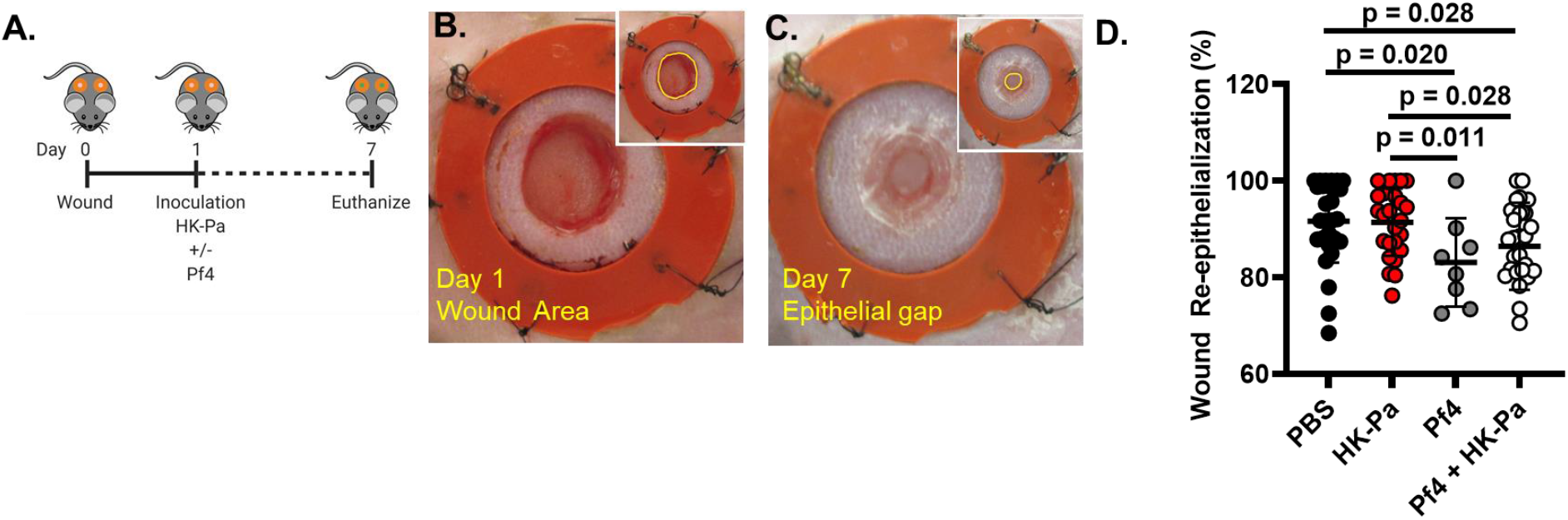
Pf phage delay re-epithelialization in the absence of bacterial infection in murine models. A) Schematic of full-thickness chronic *Pa* wound infection murine model using ring system. B) Image of a representative wound area inoculated with heat-killed *Pa* on Day 1. C) Image of the same wound area inoculated with heat-killed *Pa* on Day 7. D) Epithelial gap on Day 7 normalized to Day 1 wound area. Combined results from multiple experiments with 8-30 wounds in total per group. Statistics by unpaired Student’s T-test.

We observed that supplementation with Pf4 phage is associated with a larger epithelial gap on Day 7 when normalized to Day 1 wound area (**Figure 2D**), consistent with direct inhibition of wound healing by Pf4. Further, overall inflammatory cell counts are not altered by Pf phage supplementation (**Figure S2**). We conclude that Pf reduces re-epithelialization of murine wounds in the absence of infection. These studies indicate that Pf4 phage directly antagonizes wound healing.

### Pf phage inhibits re-epithelialization in a pig wound model

To better understand the impact of Pf4 phage on re-epithelialization we next examined whether Pf phage alters healing in a porcine burn wound model. Pig skin is anatomically and physiologically more similar to human skin, relative to rodents (Sullivan et al., 2001). Further, while mice largely rely on contraction for wound closure, both pigs and humans heal partial thickness wounds through re-epithelialization (Grillo et al., 1958; Sullivan et al., 2001). As such, the porcine wound model has been suggested to more closely mimic human disease and serve as a preferred preclinical model (Gordillo et al., 2013; Sullivan et al., 2001). Here, 8 domestic white pigs were wounded according to previously established protocols (Barki et al., 2019; Roy et al., 2014a). Each pig received 8 dorsal burn wounds which were then inoculated 3 days later with PAO1 or PAO1ΔPf4 (**Figure 3A**). At indicated times, images were taken of wounds (**Figure 3B**) and tissue healing was assessed by digital planimetry (**Figure 3B-C**). In this model Pf4 does not delay wound healing as determined by morphometric measurement over time (**Figure 3C**). The limited inflammatory change noted in *Pa* infected pig wounds (**Figure 3B**), relative to mice (**Figure 1B**) may account for these disparate results between these two animal model systems. Further, visual wound area measurements do not account for functional wound healing which requires reepithelization and establishing of barrier integrity (Wikramanayake et al., 2014).

**Figure 3.**
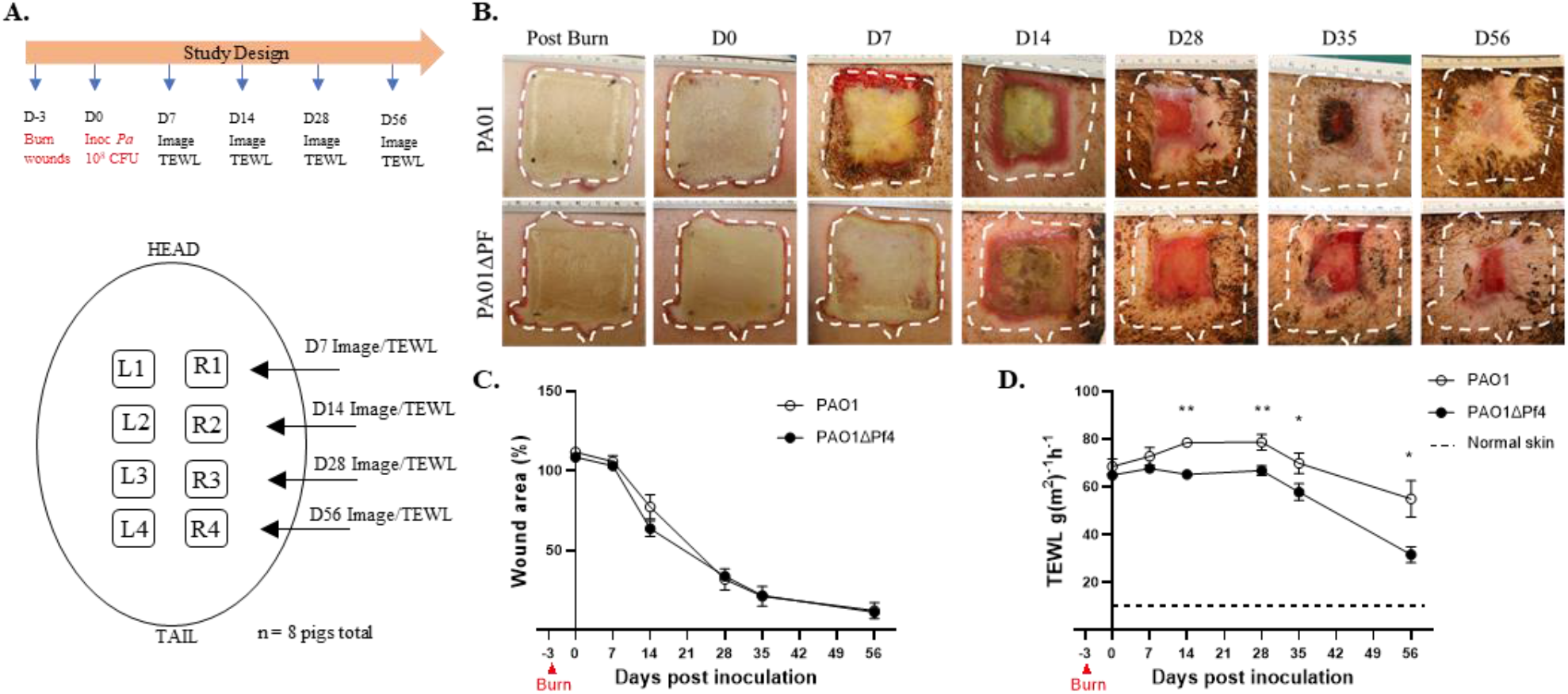
Pf is associated with increased transepidermal water loss (TEWL) in pig burn wounds. A) Schematic for porcine burn wound model with inoculation of PAO1 or PAOΔPf4. B) Representative images of wounds infected with PAO1 or PAOΔPf4 at indicated times. C) Wound area as measured by digital planimetry over time. Each independent wound represents one “n” with 32 wounds per timepoint per each treatment group. D) Wound area barrier integrity as measured by TEWL over time. Each independent wound represents one “n” for the experiment with 6-28 wounds per timepoint per treatment group. Baseline measurements done on healthy skin. *p<0.05, **p<0.01 by unpaired Student’s T test.

To assess skin barrier function integrity, following Pf4 infection, transepidermal water loss (TEWL) studies were performed on wounds at indicated time (**Figure 3A**). TEWL is a validated, objective and highly sensitive assessment of skin integrity and wound re-epithelization (Barki et al., 2019; Roy et al., 2014a). While wounds infected with PAO1ΔPf4 recover barrier function over time, and approach that of healthy skin, PAO1-infected wounds, despite apparent wound closure (**Figure 3B-C)**, demonstrate persistent water vapor loss consistent with barrier dysfunction (**Figure 3D**). These studies reveal persistent barrier dysfunction in Pf4-infected wounds, indicating that Pf phage compromises wound re-epithelization.

### Pf4 phage impair *in vitro* keratinocyte migration

The TEWL studies reveal impaired barrier integrity in Pf4-infected wounds, consistent with defects in re-epithelization. Because re-epithelialization is mediated by keratinocytes, these findings suggest Pf4 may influence keratinocyte proliferation and/or migration. Direct stimulation of keratinocytes and fibroblasts with Pf4 does not alter cell viability or proliferation (data not shown). We next assessed if Pf4 inhibits keratinocytes cell migration in an *in vitro* model commonly used to study mechanisms related to wound healing (Liang et al., 2007). A schematic of this protocol is shown (**Figure 4A)**. We demonstrate that treatment of HaCaT keratinocytes with purified Pf4 phage impairs cell migration relative to controls (**Figure 4B-C**). These effects were most pronounced in the setting of bacterial lipopolysaccharide (LPS), a component of the bacterial outer membrane that can be expected to be present wherever Pf phages are expressed by bacteria. These data suggest Pf4 phage directly antagonize keratinocyte migration in response to inflammatory signals associated with bacteria.

**Figure 4.**
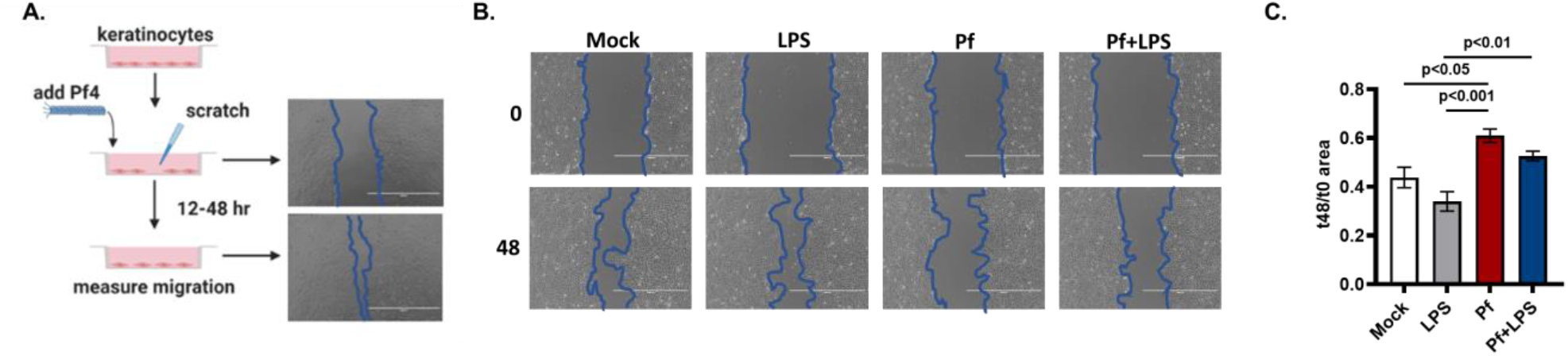
Pf phage impedes keratinocyte migration. A) Schematic keratinocyte migration assays with representative images of migration assay of HaCaT keratinocytes at 0 hour and 24 hours. B) Representative images of conditions at 0 hours and 48 hours. C) Effect of Pf4 (1 × 10^10^ copy #/ml) on HaCaT keratinocyte migration, presented as area at 24 hours/area at 0 hours with decreased area correlating with increased cell migration. Results are representative of 3 independent experiments. Statistic by one-way ANOVA.

Our previous studies revealed that Pf4 phage suppresses production of TNFα and other cytokines made in response to LPS by macrophages and monocytes (Secor et al., 2017; Sweere et al., 2019). To investigate if Pf4 alters expression of cytokines and chemokines by keratinocytes, we performed a Luminex assay on HaCaT keratinocytes treated with Pf4 phage. While Pf4 does not alter expression of the majority of cytokines and chemokines tested, select inhibition was noted of CXCL1, CCL7 and CCL22, relative to LPS-treated cells (**Figure 5A-C**). Further, treatment of HaCaT cells with Pf and LPS reduces CXCL1 and CCL22 expression relative to LPS-treated cells. (**Figure 5C**). These data argue Pf4 actively suppresses keratinocyte-derived chemokine production during inflammatory challenge and may contribute to diminished keratinocyte migration and wound re-epithelization.

**Figure 5.**
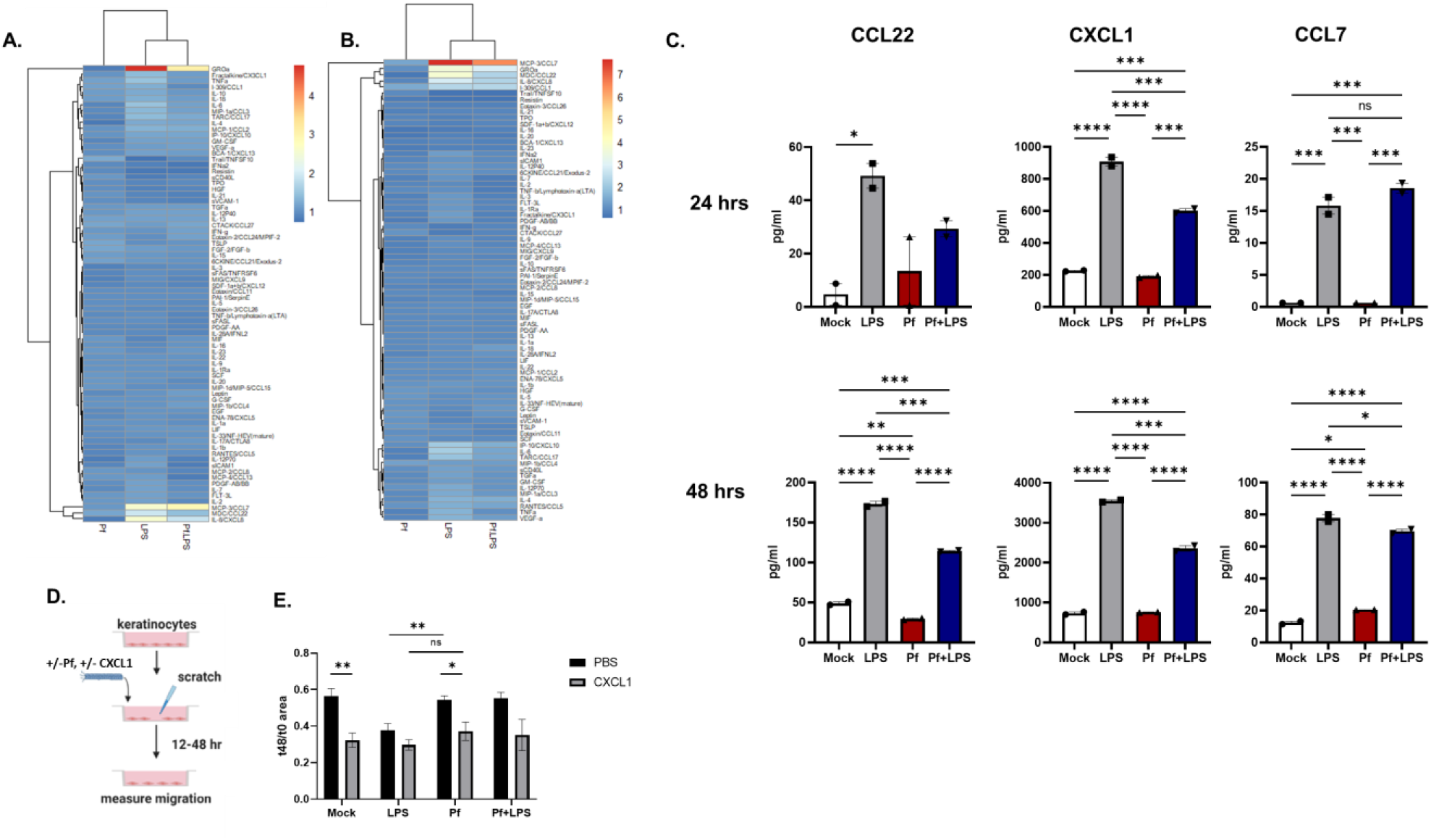
Pf inhibits keratinocyte expression of CXCL1 to inflammatory stress. Heat maps from Luminex Immunoassay performed on supernatant from HaCaT keratinocytes stimulated with Pf4 and LPS at 24 hours (A) and 48 hours (B). Results are presented as log 2 fold change of concentration compared to PBS control. C) Interpolated values (pg/ml) of select cytokines and growth factors from Luminex assay of human primary monocytes stimulated with Pf4 and LPS. *p<0.05, **p<0.01, ***p<0.001, ****p<0.0001 by one-way ANOVA. D) Schematic of keratinocyte migration assays with pretreatment with CXCL1, Pf4 and/or LPS. E) Effect of CXCL1 supplementation (1 ug/ml) on HaCaT keratinocyte migration, presented as area at 24 hours/area at 0 hours with decreased area correlating with increased cell migration *p<0.05, **p<0.01 by unpaired Student’s T test.

While CCL7 and CCL22 appear to primary recruit monocytes and lymphocytes, CXCL1 has been described to promote keratinocyte migration (Kroeze et al., 2012). To determine if Pf4-mediated CXCL1 inhibition impedes keratinocyte migration, CXCL1 or vehicle control was supplemented to keratinocytes in addition to Pf4, LPS or mock treatment. Exogenous CXCL1 did not confer additive effects to LPS treatment (**Figure 5E**). However, the addition of CXCL1 rescues keratinocyte migration among Pf4-treated cells (**Figure 5E**). Together these data indicate that Pf4 phage directly target keratinocytes and inhibit chemokine expression during inflammatory stress to limit cell migration.

### Pf phage are abundant in human wounds infected with *P. aeruginosa*

In light of the decreased wound healing observed in the presence of Pf4 phage in our *in vivo* model, we investigated the clinical impact of Pf phage on human chronic *Pa* wound infections. A total of 113 patients referred to the Infectious Disease service at the Stanford Advanced Wound Care Center (AWCC) were enrolled from June 2016 to June 2018. Our protocol for screening and classifying these patients is described in **Figure S3.** Patient characteristics are described in **Table S1.** Three *Pa-*positive patients (one who was Pf(+),and two who were Pf(-)) were excluded from the final analysis due to amputation, exposed bone/tendon, or loss to follow up (**Figure S3**). Microbiological characteristics of the *Pa*-positive wound isolates are described in **Table S2**.

Of the *Pa*-positive patients included in our analysis, Pf prophage (integrated into the bacterial chromosome) was detected in 69% (25/36) of the wounds (**Table S2**). Pf phage levels ranged from 3.55 × 10^3^ – 2.69 × 10^8^ copies/swab, with a mean of 2.16 × 10^7^ copies/swab in Pf(+) samples. Subjects in the Pf(-) subset were older compared to the Pf(+) subset (61.6 +/− 14.7 versus 76.6 +/− 14.5; p=0.008; **Table S1**). Higher numbers of Pf(+) patients were on antimicrobial treatments compared to Pf(-) patients, though this was not statistically significant (56% vs 18%; p=0.067; **Table S2**). There were no significant correlations between Pf phage status and gender, BMI, recurrence of infection, race/ethnicity, co-morbidities, antimicrobial resistance, or co-infection with other bacterial or fungal species (**Table S1**). This excludes co-infection as a confounding variable. These data indicate that Pf phage are prevalent and highly abundant in chronic wounds in this cohort.

### Pf phage are associated with wound progression

To test whether Pf phage inhibit wound healing, we followed this cohort over time for a period of two years. We had previously reported in a cross-sectional cohort study of these same patients that wounds infected with Pf(+) strains of *Pa* were significantly older at the time of enrollment than Pf(-) strains (Sweere et al., 2019). We now asked in a prospective cohort study whether wounds infected with Pf(+) *Pa* strains versus Pf(-) *Pa* strains healed over time.

A survival analysis using the endpoint of wound closure showed a significant difference between time to wound closure of the Pf(+) wounds vs Pf(-) wounds (p = 0.013 by Gehan-Breslow-Wilcoxon test) (**Figure 6A**). We further measured changes in wound dimensions over time in the 36 *Pa-*positive patients, starting with measurements taken at the patient’s initial clinic visit. We found that Pf(+) patients were more likely to experience an increase in wound size over time than patients who were Pf(-) (8/25 versus 0/11, respectively; p = 0.033 by two-sided Chi square test) (**Figure 6B-C**). We observed that all wounds that grew in size over the study period were Pf(+)(**Figure 6C**). Together, these data indicate that Pf(+) strains of *Pa* are associated with delayed wound healing.

**Figure 6.**
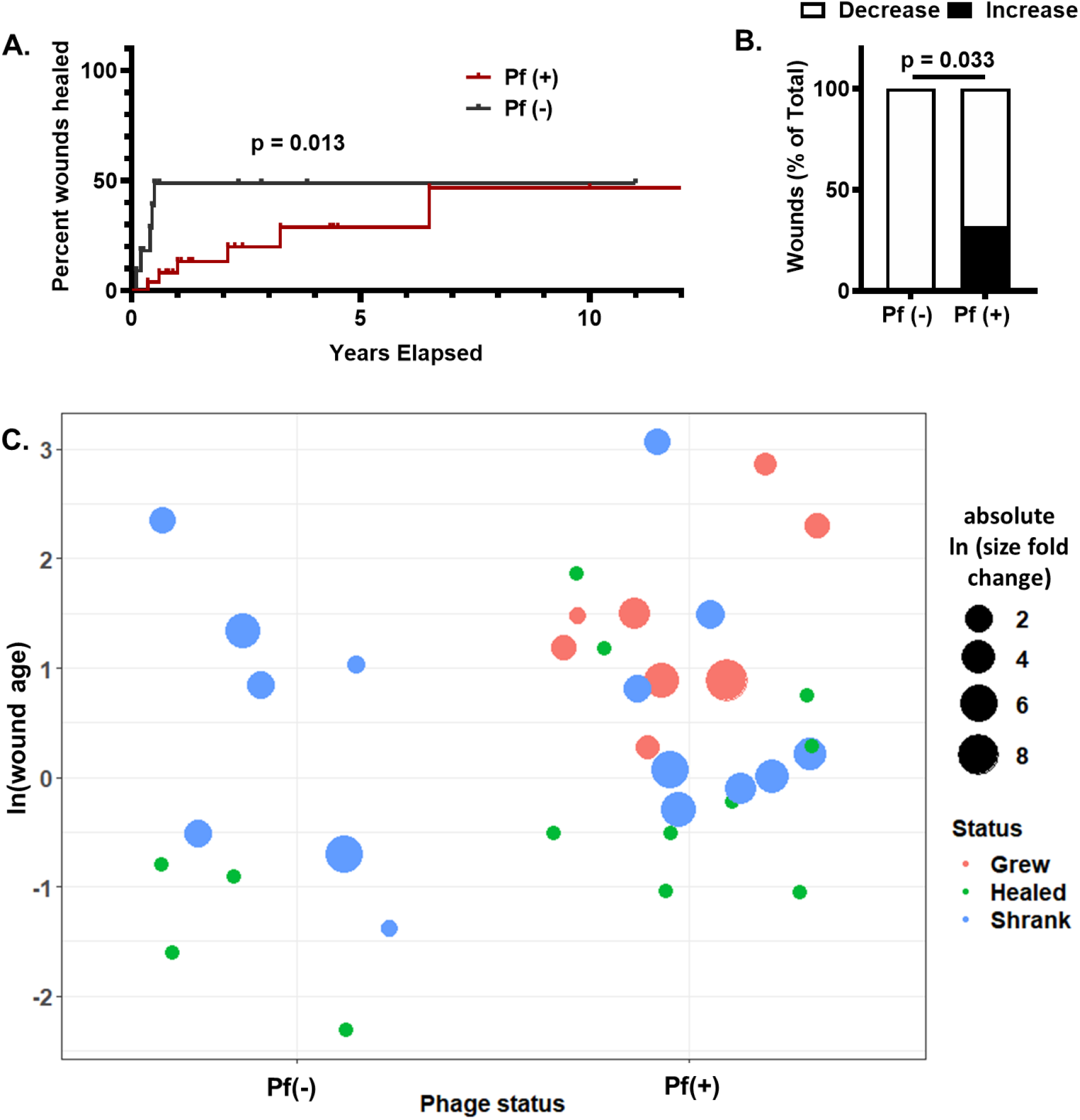
Pf phage is associated with more chronic wound infections and increased wound size in humans. A) Survival analysis of Pf(+) and Pf(-) *Pa* wound infections, with wound healing as end point. Wounds that were not healed by the end of the study were censored. p=0.013 by Gehan-Breslow-Wilcoxon test. B) Pf(+) status is associated with increases in wound size. Pf(-): n=11; Pf(+): n=25; p=0.033 by Chi square test. C) Plot showing wound age and change in size. Wounds were plotted based on Pf(-) or Pf(+) phage status and wound age (calculated as ln(wound age in years)). Wound size change is represented as absolute ln fold change, with color indicating whether wound grew (red), healed (green), or shrank (blue).

## Discussion

We report that Pf phage impairs wound healing in mice, pigs, and humans. In mice, Pf phage delays healing in both the presence or absence of live *Pa,* indicative that effects on wound healing are independent of Pf influence on *Pa* pathogenesis. In pigs, infection with a Pf(+) strain of Pa was associated with impaired re-epithelialization. Further, in a prospective cohort study in humans, Pf phage is associated with delayed wound healing and wound size progression. Together, these data strongly implicate Pf phage in delayed wound healing.

These effects of Pf phage on delayed healing are associated with altered keratinocyte chemokine expression and impaired re-epithelialization. In particular, keratinocyte secretion of CXCL1 secretion is impaired by Pf4 in the setting of inflammatory stress. Re-epithelialization is critical for wound closure and its failure leaves wounds vulnerable to reinjury, re-infection, and perpetuation (Barrientos et al., 2008;

Pastar et al., 2014; Raja et al., 2007). Our data are consistent with reports that *Pa* infections can cause impaired epithelialization (Roy et al., 2014a), although this was not previously attributed to Pf phage.

These studies align well with but are distinct from our earlier reports on Pf phages promoting chronic infection. In those studies, we reported that Pf phage inhibits bacterial clearance through effects on phagocytosis and TNF production, contributing to the establishment of chronic wound infections (Sweere et al., 2019). We likewise reported that Pf promotes chronic lung infections (Burgener et al., 2019). The data presented here indicate that in tandem with inhibiting anti-bacterial immunity, Pf phages also impair wound re-epithelialization.

Limitations of this work include that our prospective cohort study is small in size (36 patients). It will be important to validate these findings in larger, independent cohorts. It will also be important to validate these findings with Pf(+) and Pf(-) *Pa* clinical isolates as currently only lab strains were examined in our preclinical models.

This work reveals a novel insight by which bacteriophages directly modulate mammalian cells to promote disease. We conclude that Pf phage may provide a novel biomarker and therapeutic target for the delayed wound healing that occurs in the context of *Pa* wound infections. This finding may have particular relevance given our recent report that a vaccine targeting Pf could prevent initial Pa wound infection (Sweere et al., 2019). Future longitudinal studies in independent cohorts are needed to validate these studies.

## Acknowledgements

M.S.B. was supported by the Stanford Bio-X Fellowship. C.D.V. was supported by grant T32 AI007502, Doris Duke Physician Scientist Fellowship, and K08AI151089. J.M.S. was supported by the Gabilan Stanford Graduate Fellowship for Science and Engineering and the Lubert Stryer Bio-X Stanford Interdisciplinary Graduate Fellowship. P.L.B. was supported by grants R21AI133370, R21AI133240, R01AI12492093, and grants from Stanford SPARK, the Falk Medical Research Trust, and the Cystic Fibrosis Foundation (CFF).

## Author Contributions

M.S.B. contributed to the study design, conducted experiments, acquired and analyzed data, and prepared the manuscript. C.D.V. provided methods, conducted experiments, analyzed data, and revised the manuscript. AK contributed to manuscript revision. J.M.S. provided methods and assisted in conducting experiments, data analysis, and manuscript revisions. E.B.B. provided methods and assisted in data analysis. S.B. and S.K. provided methods and assisted in conducting experiments and data analysis. M.B. and V.S. assisted in conducting experiments and data analysis. P.L.B. and G.A.S. provided oversight in study design, conducting experiments, data analysis, and manuscript revision.

## Declaration of Interests

The authors declare no competing interests. G.A.S. received grants and has an equity and royalty-bearing know-how agreement with Adaptive Phage Therapeutics (APT).

## Methods

### LEAD CONTACT AND MATERIALS AVAILABILITY

- Further information and requests for resources and reagents should be directed to and will be fulfilled by the Lead Contact, Paul L. Bollyky.
- This study did not generate new unique agents.

### EXPERIMENTAL MODEL AND SUBJECT DETAILS

#### Bacterial strains and culture conditions

*P. aeruginosa* strain PAO1 was used for all experiments. Isogenic phage-free strain PAO1ΔPf4 is derived from strain PAO1 with a complete knockout of Pf4 phage entirely (Rice et al., 2009). In general, bacteria were prepared as follows. Frozen glycerol stocks were streaked on Luria-Bertani (LB) agar and grown overnight at 37°C. An isolated colony was picked and grown overnight at 37°C in LB medium, pH 7.4 under shaking, aerobic conditions. The next day, cultures were diluted to OD600 = 0.05 in 75 mL LB media and cultures were grown until early exponential phase (OD600 ± 0.3). OD600 was measured and the required number of bacteria was calculated, washed, and prepared as described in the experiments.

#### Cell Culture Conditions

HaCaT human keratinocyte cell line (AddexBio, Cat. No. T0020001) was cultured in DMEM with 10% FBS, 100 IU penicillin, and 100 ug/mL streptomycin in T-75 flasks (Corning, Cat. No. 430641). All cells were cultured on sterile tissue culture-treated plates (Falcon). Cells were cultured at 37°C, 5% CO2 at a 90% humidified atmosphere.

#### Mice

Mice were bred and maintained under specific pathogen-free conditions, with free access to food and water, in the vivarium at Stanford University. Mice that underwent surgery received additional Supplical Pet Gel (Henry Schein Animal Health, Cat. No. 029908) and intraperitoneal injections of sterile saline (Hospira, Cat. No. 0409-4888-10). All mice used for *in vivo* infection experiments were littermates. Conventional C57BL/6J mice were purchased from The Jackson Laboratory (Bar Harbor, ME). All experiments and animal use procedures were approved by the Institutional Animal Care & Use Committee at the School of Medicine at Stanford University.

#### Collection of wound swabs from human patients

From 06/2016 to 06/2018, patients visiting the Stanford Advanced Wound Care Center (AWCC) in Redwood City, California with open wounds were swabbed in duplicate over a one square inch area using Levine’s technique, using nylon-flocked wet swabs (Copan Diagnostics, Cat. No. 23-600-963). Swabs were collected in PBS and stored in −80°C before transport on dry ice. In the laboratory, the swabs in PBS were thawed, vortexed vigorously for 15 seconds, and the contents were aliquoted for quantitation of *Pa* rpIU gene and Pf prophage gene PAO717, as detailed below. Patients at the Wound Care Center were also swabbed for confirmation by diagnostic laboratory culture for the presence of *Pa.* Patients were subsequently followed until wounds completely healed or until August 2018. Patients were considered *Pa*-positive if their swabs had detectable *Pa* rpIU and their diagnostic cultures were positive. Patients were considered Pf phage-positive if both duplicate wound swabs had detectable levels of Pf phage genes. None of the *Pa*-negative patients had detectable Pf phage. For patients with multiple wounds, the dominant wound was selected for analysis. Patient enrollment and swab collection were done in compliance with the Stanford University Institutional Review Board for Human Research. Written informed consent was obtained from each patient before swab collection.

#### Human wound closure analysis

Length, width, and depth measurements of the wounds were taken and recorded into the patient’s flowsheet at the start of each visit by the intake nurse. Additional measurements of undermining and tunneling were also taken when applicable. Wound measurements were taken for all patients in the study each time they visited the Stanford Advanced Wound Care Center (AWCC). The AWCC nursing staff is trained in collecting and recording the longest length, width, and depth for each wound. Total volume was calculated for each wound for analysis. Wound healing was defined as a reduction in size compared to the first recorded measurement, whereas wound size increase was defined as an increase in wound volume compared to the first recorded measurement. Informed consent was obtained from patients before images were taken. Patient data was collected from electron medical records (EMR), including patient age, gender, co-morbidities, wound age, and other variables. This included history and physicals, progress notes, and documents uploaded into the EMR, such as the AWCC patient intake questionnaire. Patient flowsheet review was accessed for precise wound measurements and laboratory results were accessed to assess renal function and glycemic control. Microbiologic data were reviewed for antibiotic resistance profiles.

### METHOD DETAILS

#### Materials and Reagents

The following chemicals, antibiotics, and reagents were used: RPMI (HyClone, Cat. No. SH30027.01); DMEM (Hyclone, Cat. No. SH30243.01); PBS (Corning Cellgro, Cat. No. 21-040-CV); tryptone (Fluka Analytical, Cat. No. T7293); sodium chloride (Acros Organics, Cat. No. 7647-14-5); yeast extract (Boston BioProducts, Cat. No. P-950); agar (Fisher BioReagents, Cat. No. BP9744); gentamicin (Amresco, Cat. No. E737).

#### Preparation of heat-killed bacteria

Frozen glycerol stocks were streaked on LB agar as described above. Individual colonies were grown in 5 mL of LB broth the next day for 2 hours to approximately 2 × 10^8^ CFU/mL. The bacterial cultures were centrifuged at 6,000 × *g* for 5 minutes, and the pellet was washed in 1 mL of PBS three times. Finally, the pellet was resuspended in 1 mL of PBS and heated for 30 minutes at 90°C under shaking conditions. The preparation was checked for sterility by plating.

#### Phage Purification

This was performed as previously reported (Sweere et al., 2019b). In brief, phages were purified by PEG precipitation only unless noted otherwise. Bacteria were infected with stocks of Pf4 phage at mid-log phase and cultured in 75 mL of LB broth for 48 hours at 37°C under shaking conditions. Bacteria were removed by centrifugation at 6,000 × *g* for 5 minutes, and supernatant was treated with 1 μg/mL DNase I (Roche, Cat. No. 4716728001) for 2 hours at 37°C before sterilization by vacuum filtration through a 0.22 μM filter. In some experiments, supernatant was treated with 250 μg/mL of RNAse A (Thermo Fisher Scientific, Cat. No. EN0531) or 85 U/mL of benzonase (Novagen, Cat No. 70746) for 4 hours at 37°C before sterilization. Pf phage were precipitated from the supernatant by adding 0.5 M NaCl and 4% polyethylene glycol (PEG) 8000 (Milipore Sigma, Cat. No. P2139). Phage solutions were incubated overnight at 4°C. Phage were pelleted by centrifugation at 13,000 × *g* for 20 minutes, and the pellet suspended in sterile TE buffer (pH 8.0). The suspension was centrifuged for 15,000 × *g* for 20 minutes, and the supernatant was subjected to another round of PEG precipitation. The purified phage pellets were suspended in sterile PBS and dialyzed in 10-kDa molecular weight cut-off tubing (FisherScientific, Cat. No. 88243) against PBS, quantified by qPCR, diluted at least 10,000× to appropriate concentrations in sterile PBS and filter-sterilized. Three different Pf4 preparations diluted in PBS to working concentrations (1 × 10^8^ Pf4/mL) were tested for endotoxin by *Limulus* amoebocyte lysate testing at Nelson Labs (Salt Lake City, UT). All three preps had endotoxin levels under the test sensitivity level of 0.05 EU/mL.

#### Quantification of Pf phage

As several factors can produce plaques on bacterial lawns (other species of phage, pyocins, host defensins, etc.), we quantitated Pf phage using a qPCR assay as previously described (Secor et al., 2015). In brief, to quantitate Pf prophage in human or mouse wound homogenates and purified Pf phage preparations, bacterial cells and debris were removed by centrifugation at 8,000× g for 10 min. Supernatants were boiled at 100°C for 20 min to denature any phage particles, releasing intact Pf phage DNA. 2 *μL* was used as a template in 20 *μL* qPCR reactions containing 10 *μL* SensiFAST™ Probe Hi-ROX (Bioline, Cat. No. BIO-82020), 200 nM probe, and forward and reverse primers with concentrations depending on the target. Cycling conditions were as follows: 95°C 2 min, (95°C 15 sec., 60°C 20 sec.) x 40 cycles on a StepOnePlus Real-Time PCR system (Applied Biosystems). For a standard curve, the sequence targeted by the primers and probe were inserted into a pUC57 plasmid (Genewiz) and tenfold serial dilutions of the plasmid were used in the qPCR reactions. For the human wound swabs, the primers and probe were designed to recognize PAO717, a gene conserved across the Pf phage family (F at 600 nM: TTCCCGCGTGGAATGC; R at 400 nM: CGGGAAGACAGCCACCAA; probe: AACGCTGGGTCGAAG); the *Pa* 50S ribosomal gene *rpIU* was used to confirm *Pa* infection (F at 200 nM: CAAGGTCCGCATCATCAAGTT; R at 200 nM: GGCCCTGACGCTTCATGT; probe: CGCCGTCGTAAGC). For the Pf4 phage purifications, the primers and probe were designed to recognize a Pf4 phage-specific intergenic region between PAO728 and PAO729 (F at 500 nM: GGAAGCAGCGCGATGAA; R at 500 nM: GGAGCCAATCGCAAGCAA; probe: CAATTGCGCTGGTGAA). For the Pf phage purification preparations, levels of *Pa* 50S ribosomal gene *rpIU* were measured to correct for contaminating genomic *Pa* DNA, but those levels were usually negligible.

#### *In vivo* murine full-thickness wound infection model

This was done as recently described (Sweere et al., 2019a; Sweere et al., 2019b). In brief, ten-to-twelve-week old male mice were anesthetized using 3% isoflurane, and their backs were shaved using a hair clipper and further depilated using hair removal cream (Nair). The shaved area was cleaned with sterile water and disinfected twice with Betadine (Purdue Fredick Company, Cat. No. 19-065534) and 70% ethanol. Mice received 0.1-0.5 mg/kg slow-release buprenorphine (Zoopharm Pharmacy) as an analgesic. Mice received two dorsal wounds by using 6-mm biopsy punches to outline the wound area, and the epidermal and dermal layer were excised using scissors. For certain experiments, as indicated in the text, a silicone ring was sutured in place around the wound. The wound area was washed with saline and covered with Tegaderm (3M, Cat. No. 1642W). Bacteria were grown as described above and diluted to 1 × 10^7^ CFU/mL in PBS. Mice were inoculated with 40 μL per wound 24 hours post-wounding, and control mice were inoculated with sterile PBS. Mice were weighed daily and given Supplical Pet Gel and intraperitoneal injections of sterile saline. Upon takedown, wound beds were excised and processed for histological analysis.

#### Wound Healing Analysis in Murine Models

This was done as previously described (Balaji et al., 2014). In brief, using a dot ruler for standardization, images of both wounds for all mice were taken on each day from Day 1 to Day 13 post-wounding. Images were taken using Canon PowerShot Camera (Item Model No. 1096C001) mounted on a fixed tripod for consistency in the height and angle at which images were taken. Using the images, wound area was measured using Fiji (Image J, National Institutes of Health) (Schindelin et al., 2012) by tracing the border of the wounded area, as well as the border of a standardized dot measure. For histologic analyses, wounds were harvested, bifurcated, fixed in 10% neutral buffered formalin, and embedded in paraffin. Wound sections of 5 μm were cut from paraffin-embedded blocks. Immune cell influx, general tissue morphology, epithelial gap and granulation tissue area were measured from hematoxylin and eosin (H&E) stained sections using morphometric image analysis (Image-Pro, Media Cybernetics, Silver Spring, MD, and Metamorph, Molecular Devices, Downingtown, PA). Images were obtained using a Nikon Eclipse microscope, and image analysis was performed using the Nikon Elements software (Nikon Instruments, Melville, NY, USA). The percentage of CD45+ cells to the total cell infiltrate was calculated in 6 high-powered fields (HPFs, 64X) for each wound section. The HPFs were chosen just above the panniculus carnosus and were evenly distributed between the two wound epithelial margins. On a 4× edge-to-edge wound section image, epithelial gap was measured as the distance (in millimeters) between encroaching epithelial margins. The granulation tissue area was measured (in mm^2^) as the entire cellular region within the epithelial margin. Any sample that did not yield reliable counting due to sample quality was excluded from the analysis for that particular variable. All scoring was done by an independent pathologist.

#### Porcine Full Thickness Burn and Biofilm Wound Model

Domestic Yorkshire/landrace cross female pigs were wounded and infected to establish chronic wound biofilm model as described previously (Barki et al., 2019; Roy et al., 2014b, 2020). Eight full thickness burn wounds (2 x 2 inch) were made on the dorsum of pigs. On day 3 post-burn, wounds were inoculated with either PAO1, or PAO1ΔPf4 strains (CFU10^8^/ml). Wounds were followed up to 56 days post inoculation. Wounds were dressed with Tegaderm ^TM^ (3M) which was kept in place with V.A.C. drape (Owens & Minor) and then wrapped with Vetrap^TM^ and Elastikon™ (3M). Dressings were changed weekly for the duration of the study. Digital images and trans-epidermal water loss (TEWL) was collected on day 0, day 7, day 14 and day 56 post inoculation. Biopsies were collected on days 7, 14, 28, and 56 post infection. The pigs were euthanized at day 56 post inoculation. Biopsies were collected using a 6 mm sterile disposable punch biopsy tool. Each independent porcine wound is considered as one “n” for the porcine wound experiments. A total of 8 pigs were used for this study.

#### Trans-epidermal Water Loss (TEWL) measurement

DermaLab Combo™ (cyberDERM inc., Broomall, PA) was used to measure the trans-epidermal water loss from the wounds. TEWL was measured in g (m^2^)^-1^ h^-1^ (Roy et al., 2014b). Dermalab Combo consists of a main measuring unit with a computer and a probe. The TEWL probe has two hygro sensors located close to each other in a perpendicular orientation, TEWL is determined from the humidity gradient between the sensors. The probe is placed on the porcine skin/wound surface. The evaporated water released from the skin/wound is detected by the sensors in the probe and measured to provide the TEWL value.

#### Migration Assay

For HaCaT cell migration assays a standard scratch assay was used (Cory, 2011). In brief, cells were seeded at a density of 1×10^6^ cells per well in DMEM 10% FBS in collagen-coated 6-well tissue culture plates. Media was removed and HaCaT cells were then serum starved in DMEM containing only 2% FBS for 24 hours prior to the treatment. A scratch defect was created in the cell monolayer along the diameter in each group using a sterile pipette tip (200 μl tip). Treatment groups included Pf4 bacteriophage with final concentration of 1 × 10^10^ copy #/ml and/or LPS 1 ug/ml. For supplementation studies, HaCaT cells were treated with CXCL1 (Peprotech, Cat no 250-11) at a final concentration of 1ug/ml in addition to Pf4 or LPS described as above. Photographic images were obtained at 0 and 24 hours incubation using an Invitrogen EVOS FL Imaging System. The unfilled scratch defect area was measured at each reference point per well. Data was presented as extent of wound closure, that is, the percentage by which the scratch area has decreased at a given time point for each treatment as compared to the original defect (at 0 hours). All experiments were carried out at minimum in triplicate and the passage number was similar amongst the different groups.

#### Luminex immunoassay

HaCaT cells were seeded at 1×10^6^ cells per well in DMEM 10% FBS in collagen-coated 6-well tissue culture plates. Media was removed and HaCaT cells were then serum starved in DMEM containing only 2% FBS for 24 hours prior to the treatment. Cells were treated with Pf4 (1×10^10^ copies/ml) and/or LPS (1 μg/ml). Supernatant was collected at 24 and 48hrs and used for cytokine profiling through Luminex (The Human Immune Monitoring Center, Stanford). Human 63-plex kits were purchased from eBiosciences/Affymetrix and used according to the manufacturer’s recommendations with modifications as described below. Briefly: Beads were added to a 96 well plate and washed in a Biotek ELx405 washer. Samples were added to the plate containing the mixed antibody-linked beads and incubated at room temperature for 1 hour followed by overnight incubation at 4°C on an orbital shaker at 500-600 rpm. Following the overnight incubation plates were washed in a Biotek ELx405 washer and then biotinylated detection antibody added for 75 minutes at room temperature with shaking. Plate was washed as above and streptavidin-PE was added. After incubation for 30 minutes at room temperature wash was performed as above and reading buffer was added to the wells. Each sample was measured in duplicate. Plates were read using a Luminex 200 instrument with a lower bound of 50 beads per sample per cytokine. Custom assay Control beads by Radix Biosolutions are added to all wells.

### QUANTIFICATION AND STATISTICAL ANALYSIS

All statistical analyses were done using GraphPad Prism (GraphPad Software, Inc. La Jolla, CA). All Unpaired T-Tests, Mann-Whitney Test, Fisher’s Exact Tests, and Chi-Square Test were two-tailed. Depicted are Means with Standard Error or Standard Deviation of the population unless otherwise stated. Statistical significance was considered *p*<0.05.

### DATA AND CODE AVAILABILITY

Original/source data for figures and tables in this paper is available by request

## Notes

### Competing Interest Statement

The authors have declared no competing interest.

