## Supplemental Figures for "Filamentous Bacteriophage Delay Healing of Pseudomonas-Infected Wounds"

| | Pa(+)Pf(+) | Pa(+)Pf(-) | p-value |
|---|---|---|---|
| Samples (n, %) | 25, 69% | 11, 31% | - |
| **Age of Patients (years)** | | | |
| **Mean (SD)** | 61.6 (14.7) | 76.6 (14.5) | 0.008 |
| Gender (n) | | | |
| Male | 9 (36%) | 6 (55%) | 0.465 |
| Female | 16 (64%) | 6 (45%) | |
| Race/Ethnicity | | | |
| Caucasian | 17 (68%) | 5 (45%) | 0.144 |
| Asian | 3 (12%) | 1 (9%) | |
| Hispanic | 3 (12%) | 5 (45%) | |
| African-American | 2 (8%) | 0 (0%) | |
| Body Mass Index (kg/$m^2$) | | | |
| Mean (SD) | 27.95 (11.9) | 30.28 (6.9) | 0.551 |
| **Age of Wound (years)** | | | |
| **Median** | 2.1 | 0.5 | 0.046 |
| Recurrence of Infection | 9 (43%) | 3 (43%) | 0.756 |
| Co-Morbidities | | | |
| Diabetes Mellitus | 12 (48%) | 3 (27%) | 0.295 |
| Renal Disease | 8 (32%) | 1 (9%) | 0.223 |

**Table S1. Patient Demographic Data.** Clinical information on patients from the AWCC with culture-positive and qPCR-positive *Pa*-infected non-healing wounds who participated in the wound swab study. Renal Disease was defined as patients with Chronic Kidney Disease (CKD) or End Stage Renal Disease (ESRD). Statistical significance was measured using Fisher's Exact Test for the following parameters: Gender, Infection Recurrence, and Co-Morbidities. Statistical significance was measured using an Unpaired T-Test for Age and BMI, Chi-Square Test for Race/Ethnicity, and Unpaired Two-Tailed Mann-Whitney Test for Age of Wound because the data was nonparametric. 9 patients were excluded from the Recurrence of Infection analysis because they did not provide an answer on the patient intake form provided by the AWCC.

| | Pa(+)Pf(+) | Pa(+)Pf(-) | p-value |
|---|---|---|---|
| Pf Phage, copies/swab, mean (range) | $2.16 \times 10^7$ ($3.55 \times 10^3$ - $2.69 \times 10^8$) | 0 | - |
| *P. aeruginosa*, copies/swab, mean (range) | $7.55 \times 10^7$ ($6.82 \times 10^2$ – $9.2 \times 10^8$) | $2.74 \times 10^7$ ($5.56 \times 10^3$ – $1.79 \times 10^8$) | 0.395 |
| Antimicrobials at Time of Swab | | | |
| **Antimicrobials** | 14 (56%) | 2 (18%) | 0.067 |
| Anti-Pseudomonals | 6 (24%) | 2 (18%) | >0.999 |
| Antibiotic Resistance | | | |
| Levofloxacin | 8 (32%) | 1 (9%) | 0.223 |
| Ciprofloxacin | 5 (20%) | 1 (9%) | 0.643 |
| Imipenem | 4 (16%) | 4 (36%) | 0.214 |
| Meropenem | 4 (16%) | 3 (27%) | 0.650 |
| Gentamicin | 2 (8%) | 1 (9%) | >0.999 |
| Piperacillin | 1 (4%) | 1 (9%) | 0.524 |
| Co-Infection | | | |
| Presence of Co-Infection | 19 (76%) | 7 (64%) | 0.454 |
| *Staphylococcus aureus* | 10 (40%) | 4 (36%) | >0.999 |
| Gram-Positive | 8 (32%) | 2 (18%) | 0.690 |
| Other Gram-Negative | 11 (44%) | 5 (45%) | >0.999 |
| *Candida* Species | 0 (0%) | 1 (9%) | 0.306 |

**Table S2. Human Wound Microbiology Data.** Statistical significance for *Pa* copies/swab was analyzed using an Unpaired T-Test. Statistical significance for Antimicrobials at Time of Swab, Antibiotic Resistance, and Co-Infection was analyzed using Fisher's Exact Test. Anti-Pseudomonal antibiotics included in the analysis were ciprofloxacin and levofloxacin. 0 patients were resistant to Tobramycin. None of the Co-Infection parameters were significant, which strengthens the correlation between Pf phage and clinical outcomes.

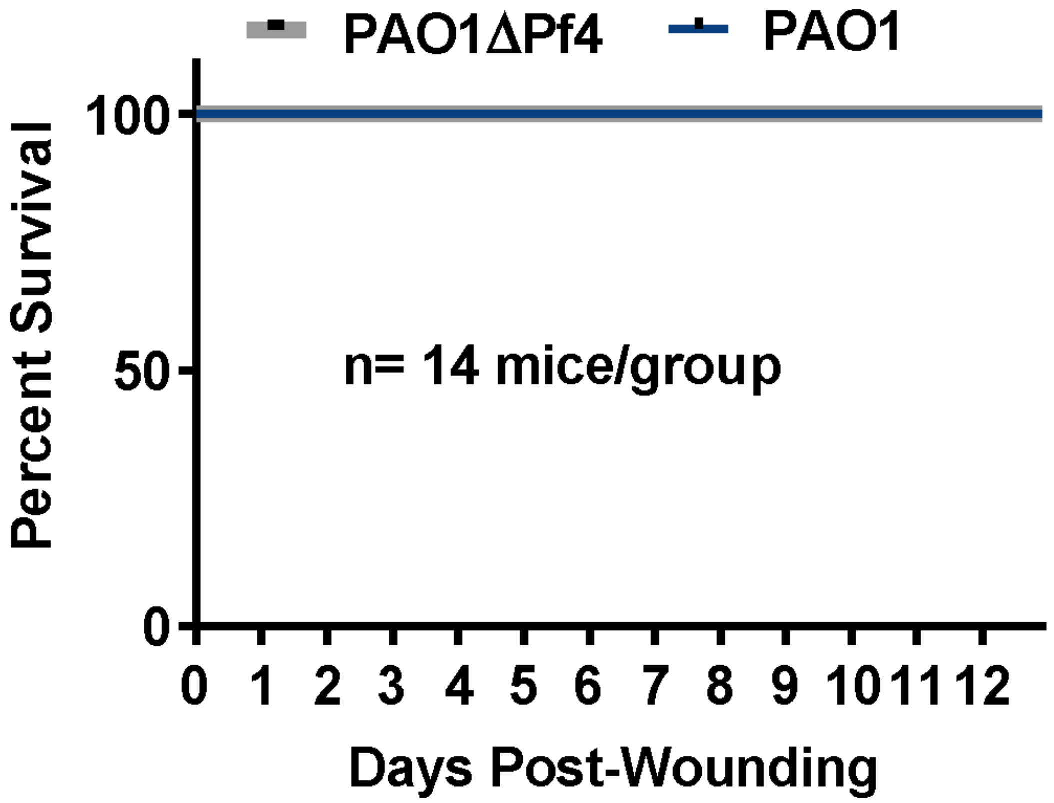


**Supplemental Figure S1. Percent Survival of Mice.** Kaplan-Meier plot of survival of mice infected with either PAO1 or PAO1ΔPf4 using the chronic *Pa* wound infection model. All mice survived until Day 13 of the experiment, when they were euthanized.

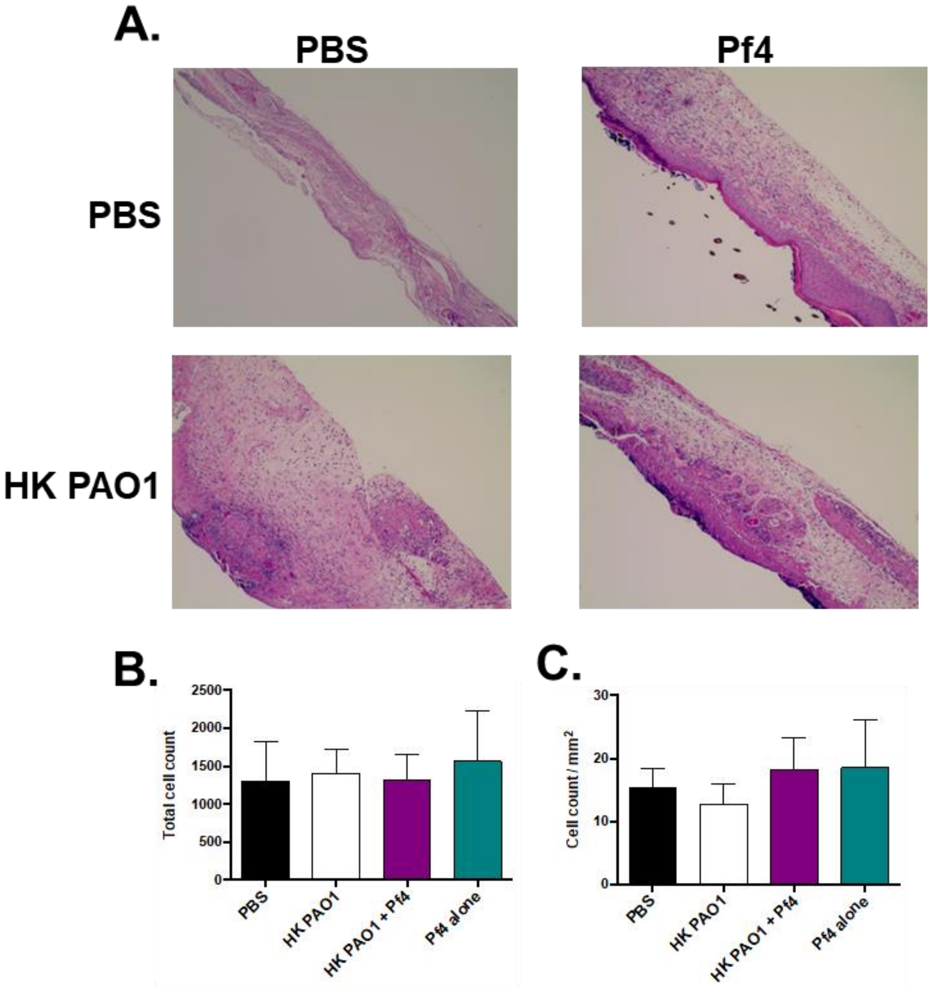


**Supplemental Figure S2: Pf phage do not alter wound cell counts in murine models.**
A) Representative H&E stains of formalin-fixed, paraffin-embedded samples from mouse wounds inoculated with heat-killed *Pa* (HK-PAO1) and/or Pf4. B) Total cell count based on H&E staining. C) Cell count/mm$^2$ based on H&E staining.

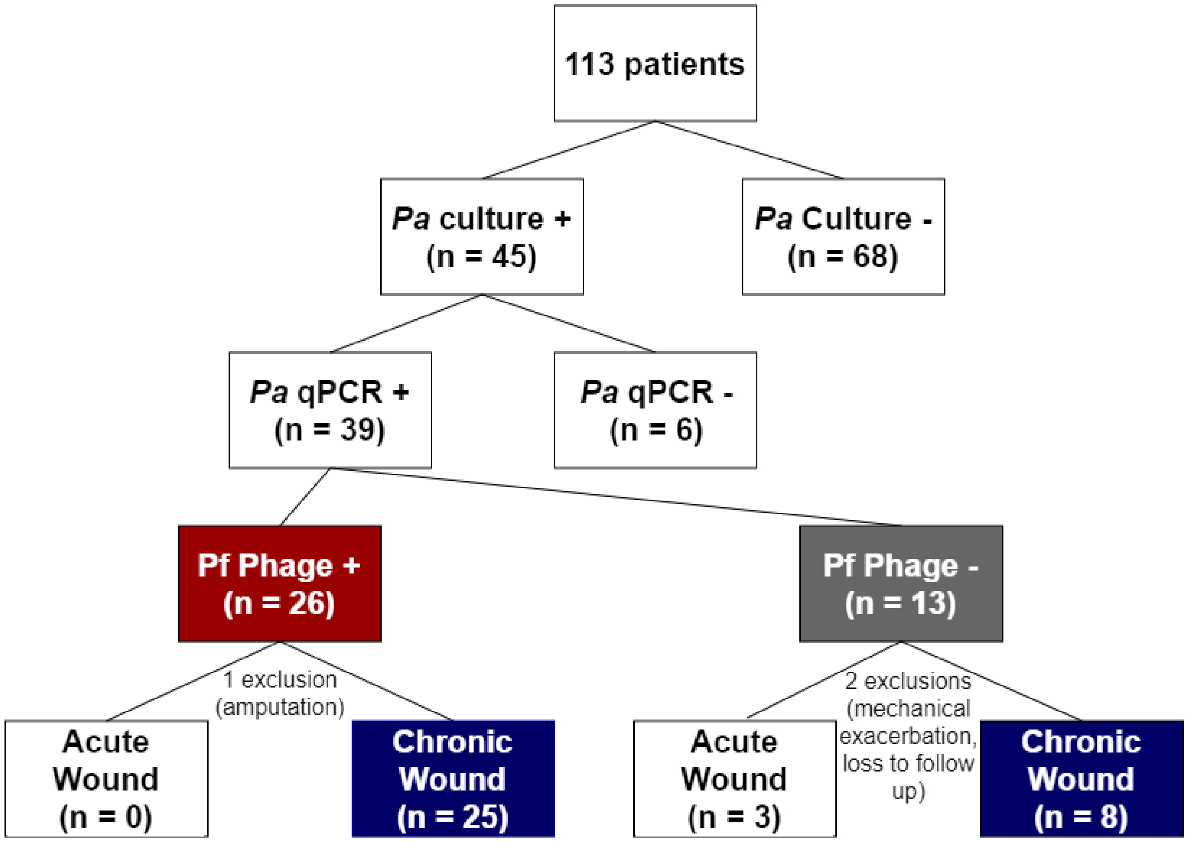


**Supplemental Figure S3.** Epidemiological flow chart. A total of 113 patients from the Stanford Advanced Wound Care Center (AWCC) were enrolled in a prospective cohort study. 39 patients were classified as Pa-positive based on culture and qPCR results. 26 patients were classified as Pf phage-positive based on qPCR results. 1 Pf phage-positive patient was excluded from the study due to an amputation (see Figure S5A). 13 patients were classified as Pf phage-negative based on qPCR results. 1 Pf phage-negative patient was excluded from the study due to a mechanical exacerbation (see Figure S5B). 1 Pf phage-negative patient was excluded from the study due to loss to follow up. Acute wounds are defined as equal to or younger than 3 months of age. Chronic wounds are defined as older than 3 months of age.
